# The Genetic Immune Basis of Gout: Identification, Functional Characterization, and Therapeutic Implications

**DOI:** 10.1101/2025.09.12.675974

**Authors:** Ali Alamdar Shah Syed, Aadil Ahmed Memon, Aamir Fahira, Qiangzhen Yang, Manfei Zhang, Xuemin Jian, Yongyong Shi

## Abstract

Gout, the most common inflammatory arthritis, is still managed mainly as a metabolic disorder, with treatment centered around urate lowering rather than blocking the immune response that drives gout flares. This emphasis reflects a historical view of gout as a disease of excess and diet, reinforced by genome-wide association studies (GWAS) that consistently detect urate transport and metabolism loci, while immune-related loci have failed to reach genome-wide significance. Growing clinical and experimental evidence, however, points to a central role for immune pathways in gout pathogenesis.

We set out isolate the immune genetic component of gout. Using a conjunctional false discovery rate (conjFDR) framework, we integrated gout GWAS data with those from eight immune-mediated disorders rheumatoid arthritis, Crohn’s disease, inflammatory bowel disease, psoriasis, multiple sclerosis, asthma, chronic obstructive pulmonary disease, and systemic lupus erythematosus. We observed robust pleiotropic enrichment across all comparisons and mapped the resulting loci through FUMA to 85 unique credible genes broadly distributed across the genome, with no classical urate transporters present. Additionally, we identified 16 novel genes for gout, many of which are of obvious immune nature.

We performed GO and KEGG enrichment analyses, which at a stringent q-value threshold identified adaptive and innate immune pathways, including T and B cell receptor signaling, antigen processing and presentation, NF-κB signaling, and Jak-STAT signaling. At a nominal p-value threshold, we uncovered additional cytokine-driven processes such as *IL6* and *IL7* signaling. We confirmed disease relevance with DisGeNET and established causality for 14 genes through Mendelian randomization, including *IL1RN, MAP3K11*, and *SH2B3* genes with existing pharmacological inhibitors.

We are the first to genetically isolate the genetic immune component of gout. Our findings show that these immune pathways can be specifically targeted, and immune medication should be incorporated into therapeutic strategies to complement urate-lowering approaches.

## Introduction

Gout is a type of inflammatory disorder characterized by the deposition of monosodium urate (MSU) crystals causing a red, tender, hot swelling of joints and soft tissues (Dalbeth et al., 2019). Uric acid is the final product of purine metabolism, under physiologic conditions it circulates as the deprotonated urate anion combining with sodium ions to form MSU(Li et al., 2023). Hyperuricemia, a condition where uric acid levels in the blood exceed 6.6 mg/dL, may over time lead to MSU depositions into joints and surrounding tissues which can trigger inflammation and a subsequent immune response that can cause significant discomfort and pain leading to disability. The prevalence of gout is reported to be 2–6-fold higher in men compared to women, while most alarmingly the global incidence of gout has increased by 63.44% over the past 2 decades representing a challenge to health and wellbeing globally (Asghari et al., 2024).

Traditionally gout has been thought to be caused by dietary intake with excess of alcohol and meat, particularly sea food and this is partly supported by studies of cohorts over decades that showed that meat and alcohol consumption increased the risk of gout in dose dependent manner (Choi, Atkinson, Karlson, Willett, & Curhan, 2004; Choi & Curhan, 2007). Therefore, current guidelines followed by clinicians mainly recommend abstaining from alcohol, reduction in meat consumption and maintenance of a healthy weight as key to reducing the incidence of gout flares (Nielsen, Zobbe, Kristensen, & Christensen, 2018; Sautner, Eichbauer-Sturm, Gruber, Lunzer, & Puchner, 2022). Modern treatment of gout revolves around the reduction and management of serum urate levels, with drugs like Allopurinol and Febuxostat being commonly prescribed urate lowering therapies. In addition to the aforementioned drugs, Colchicine is also routinely prescribed to gout patients, even though its issues with toxicity are well known(Dalbeth et al., 2019; Finkelstein et al., 2010; FitzGerald et al., 2020). However, results from mendelian randomization analysis have shown that alcohol consumption does not causally affect the risk of developing gout, showing that gout is indeed a complex disorder whose cause warrants further study (Syed et al., 2022).

We hypothesize that gout is a result of the complex interplay between products of metabolism and inflammatory response over decades of questionable lifestyle choices and that hyperuricemia should be seen as a prerequisite to gout but not the actual cause, while the individual susceptibility to gout is mainly driven by immune factors. This is supported by recent longitudinal studies which have found that the relationship between gout and serum uric acid levels is non-linear and only about half of individuals with serum uric acid levels of >10 mg/dL develop gout and not all those who experience gout flares even have visible MSU deposits in their joints (Bongartz et al., 2015; Dalbeth et al., 2018). In individuals that develop gout the immune response to the MSU crystals is led by an inflammasome known as NLRP3 (NOD-, LRR- and pyrin domain-containing 3 protein), part of innate immune system, that comprises of multiple proteins that join together to form a complex. Monosodium-urate crystals initiate the signal which causes NLRP3 to assemble and activate and form a complex cascade that involves multiple proinflammatory molecules, such as *IL1β* that will recruit neutrophils the affected joints. Neutrophils then drive the localized immune response, which will eventually be resolved by immune factors such as *IL-1ra* (*IL1RN*), *IL-37, IL-10*, and *TGF-β* (Dalbeth, Gosling, Gaffo, & Abhishek, 2021).

The fact that *IL-1* inhibitors like Anakinra were first trialed in acute gout nearly two decades ago yet remain underutilized in routine clinical practice reflects a persistent perception of gout as primarily a metabolic disorder (due to hyperuricemia) rather than an inflammatory or autoimmune-like condition (driven by NLRP3 inflammasome activation and *IL1β* mediated inflammation) (So, 2019). The slow adoption highlights a broader disconnect, while gout’s pathophysiology is increasingly recognized as auto-inflammatory (An, Marwaha, & Laxer, 2024; Galozzi, Bindoli, Doria, Oliviero, & Sfriso, 2021; Punzi, Scanu, Ramonda, & Oliviero, 2012), its clinical management remains rooted in metabolic correction. This may be partly due to the fact that for decades now genetic studies of gout have predominantly identified loci associated with urate transporters, with little implication of immune-related genes to date (Dalbeth, Stamp, & Merriman, 2017; Major, Dalbeth, Stahl, & Merriman, 2018; Reginato, Mount, Yang, & Choi, 2012). Although the scientific community is increasingly more open to recognizing gout as an autoimmune disorder rather than a purely metabolic one, characterizing its immune component has been challenging, likely due to the high degree of heterogeneity within the genetic immune component of gout compounded by the additional challenge of disentangling the genetics of gout from that of hyperuricemia.

In this study, we aim to elucidate the immune component of gout by integrating summary-level statistics from other immune disorders and identifying their overlap with gout using Conjunctional False Discovery Rate (conjFDR) analysis. We hypothesize that if gout has a genetic immune component, it may share genetic loci with other immune-related disorders. By integrating overlapping elements between gout and these disorders, we seek to compile the immune component of gout. Additionally, we aim to identify potential therapeutic targets within this immune component through Mendelian randomization and to explore the possibility of repurposing existing drugs for gout treatment.

## Material and Methods

### Study Design and Rationale

To identify the immune-related genetic component of gout, we hypothesize that gout shares a common immune component with other immune disorders. We selected eight immune disorders, Rheumatoid Arthritis, Crohn’s disease, Psoriasis, Inflammatory Bowel Disorder, Asthma, Systemic Lupus Erythematosus, Multiple Sclerosis, and Chronic Obstructive Pulmonary Disorder and examined the shared genetic loci between gout and each disorder. By aggregating these overlapping loci across multiple disorders, we aim to assemble the immune component of gout.

### Determining the genetic overlap between gout and other immune disorders

In this study, we utilized GWAS datasets for gout and eight immune-related disorders. For gout, we used the largest available dataset with 763,813 individuals (13,179 cases) from the UK Biobank (Tin et al., 2019). Rheumatoid Arthritis (RA) data were sourced from a meta-analysis of 276,020 individuals across five ancestral groups by Ishigaki et al. (Ishigaki et al., 2022). Systemic Lupus Erythematosus (SLE) data came from a study by Sakaue et al., comprising 659,165 samples (Sakaue et al., 2021). Asthma data were extracted from a UK Biobank cohort study by Valette et al., with 408,442 samples (Valette et al., 2021). Psoriasis data were from a study by Stuart et al., involving 44,161 samples(Stuart et al., 2015). Multiple Sclerosis (MS) data were from a 2011 study by the International Multiple Sclerosis Genetics Consortium led by Stephen Sawcer, with 38,135 samples (Sawcer et al., 2011). Inflammatory Bowel Disease (IBD) and Crohn’s Disease data were from a study by de Lange et al., with sample sizes of 59,957 and 40,266 individuals, respectively(de Lange et al., 2017). Lastly, Chronic Obstructive Pulmonary Disease (COPD) data were from a study by Kim et al., using the subset of never-smokers (129,175 individuals) to enrich for the immune component of the disease (Kim et al., 2021).

To identify genomic loci shared between gout and other immune disorders, we utilized the conjFDR method proposed by Andreassen et al. in 2013 (Andreassen et al., 2013). *P*-values were transformed to the −log10(*p*) scale for enhanced interpretability and visualization, commonly used in conditional Manhattan plots. Linkage disequilibrium (LD) pruning was applied to prevent correlated SNPs from inflating significance levels artificially. An FDR threshold of <0.05 was used to determine significant SNP associations. This method enables the detection of pleiotropic associations, contributing to the understanding of shared genetic architecture between gout and other diseases.

### Annotation of genes and characterization of the immune component

To characterize the significant genetic loci identified in the conjFDR analysis, we employed the FUMA pipeline function GENE2FUNC (Watanabe, Taskesen, van Bochoven, & Posthuma, 2017), a widely recognized tool for functional mapping and annotation. To prepare the conjFDR results for FUMA analysis, we used the fdr_fuma_snp_input.sh script from the *yunhoop* repository (GitHub: precimed/yunhoop), which transformed the conjFDR output into a format compatible with FUMA. Within GENE2FUNC, we performed GO and KEGG enrichment analyses to assess overrepresentation of biological processes, molecular functions, and signaling pathways among the identified genes (“The Gene Ontology resource: enriching a GOld mine,” 2021; Kanehisa, Furumichi, Sato, Kawashima, & Ishiguro-Watanabe, 2023). To complement pathway enrichment, we systematically queried the gene set against DisGeNET v7.0, a curated database of gene–disease associations, to identify links with immune-mediated and inflammatory disorders (Piñero et al., 2020).

### Mendelian randomization and drug repurposing

In this thesis we conducted two-sample Mendelian randomization (MR) in R using the TwoSampleMR package, treating genetically predicted expression of each of the 85 immune-component genes as the exposure and gout as the outcome (Tin et al., 2019); exposure instruments were derived from the eQTLGen blood consortium (Võsa et al., 2021), selecting significant cis-eQTL SNPs, pruning for independence (LD r^2^ < 0.01), and harmonizing exposure and outcome summary statistics to the same effect allele with strand-ambiguous variants excluded. The primary estimator was inverse-variance weighted (IVW), with single-SNP instruments analyzed via the Wald ratio; potential directional pleiotropy was assessed using the MR–Egger intercept, and robustness was evaluated with the weighted-median method (and leave-one-out checks where relevant). Unless otherwise stated, hypothesis tests were evaluated at α = 0.05 given the exploratory nature of this gene-set analysis. Potential target genes were used to query DGIdb and ConnectivityMap databases to identify potential drugs that could be repurposed for gout.

## Results

### Shared Genetic Loci between Gout and Immune Disorders

When using conjFDR to determine shared genetic loci between gout and eight immune-mediated disorders, we identified a substantial number of pleiotropic associations across all comparisons. After filtering for independence, the number of shared loci varied by trait, with overlaps detected between gout and Crohn’s disease (19 loci), rheumatoid arthritis (9), inflammatory bowel disease (8), psoriasis (7), multiple sclerosis (7), asthma (5), chronic obstructive pulmonary disease (4), and systemic lupus erythematosus (2). These loci were distributed across multiple chromosomes, including chromosomes 1, 2, 3, 4, 5, 6, 10, 11, 12, and 19, indicating that the genetic sharing between gout and immune-mediated disorders is not confined to a single genomic region but spread broadly across the genome.

The quantile–quantile (QQ) plots of nominal versus expected –log10 P values (*Figure 1*) illustrate the extent of pleiotropic enrichment for gout associations conditional on each immune disorder. All eight trait pairs showed a clear upward deviation from the null expectation, indicating enrichment of true pleiotropic signals. The magnitude of this deviation varied, with the largest shifts observed in Crohn’s disease, rheumatoid arthritis, IBD, psoriasis, and MS, and more moderate yet evident deviations for asthma, COPD, and SLE. Importantly, the initial portions of the QQ curves closely followed the diagonal under the null, with no early inflation, suggesting that the observed enrichment patterns are not artefacts of population stratification or sample overlap but instead reflect genuine shared genetic architecture. Across all comparisons, the consistent departure from the diagonal expectation line at lower P values supports the presence of true polygenic overlaps beyond chance.

**Figure 1.**
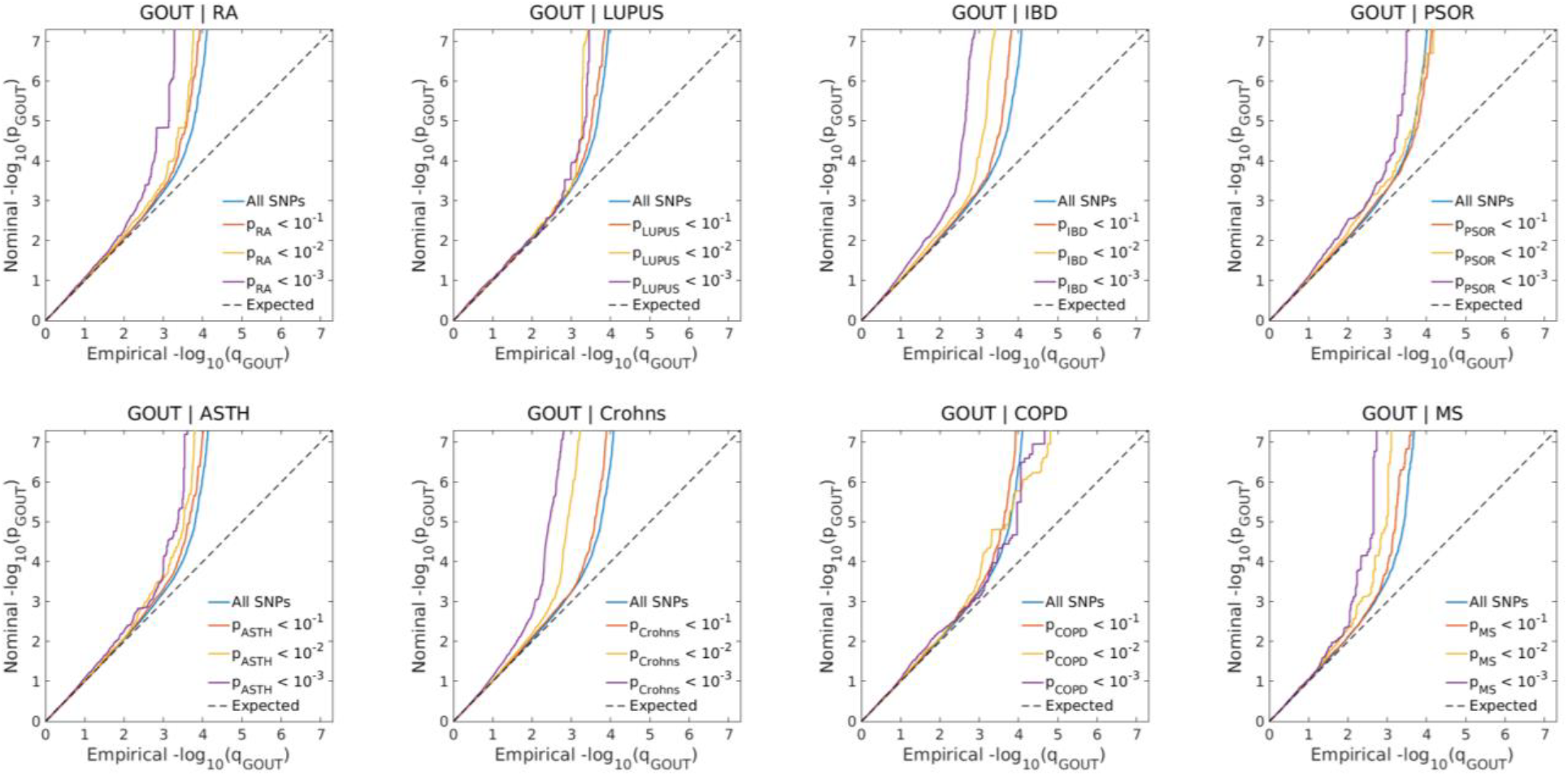
Conditional QQ plots showing nominal versus empirical –log_10_ P values for gout when conditioned on different significance thresholds in eight immune-mediated disorders. Lines represent all SNPs (blue) and SNP subsets meeting increasing significance in the conditioning trait (orange, yellow, purple). The dashed line shows the null expectation.

The conditional true discovery rate (TDR) plots (*Figure 2*) provide a complementary perspective by showing how the estimated proportion of true discoveries for gout changes when SNPs are stratified by their significance in the conditioning immune trait. In each disorder, more stringent conditioning thresholds yielded higher TDR values, reflecting increased confidence in the pleiotropic associations. The steepness and magnitude of TDR gains varied by trait, with Crohn’s disease, RA, IBD, psoriasis, and MS showing substantial enrichment patterns, while asthma, COPD, and SLE displayed smaller but still measurable improvements.

**Figure 2.**
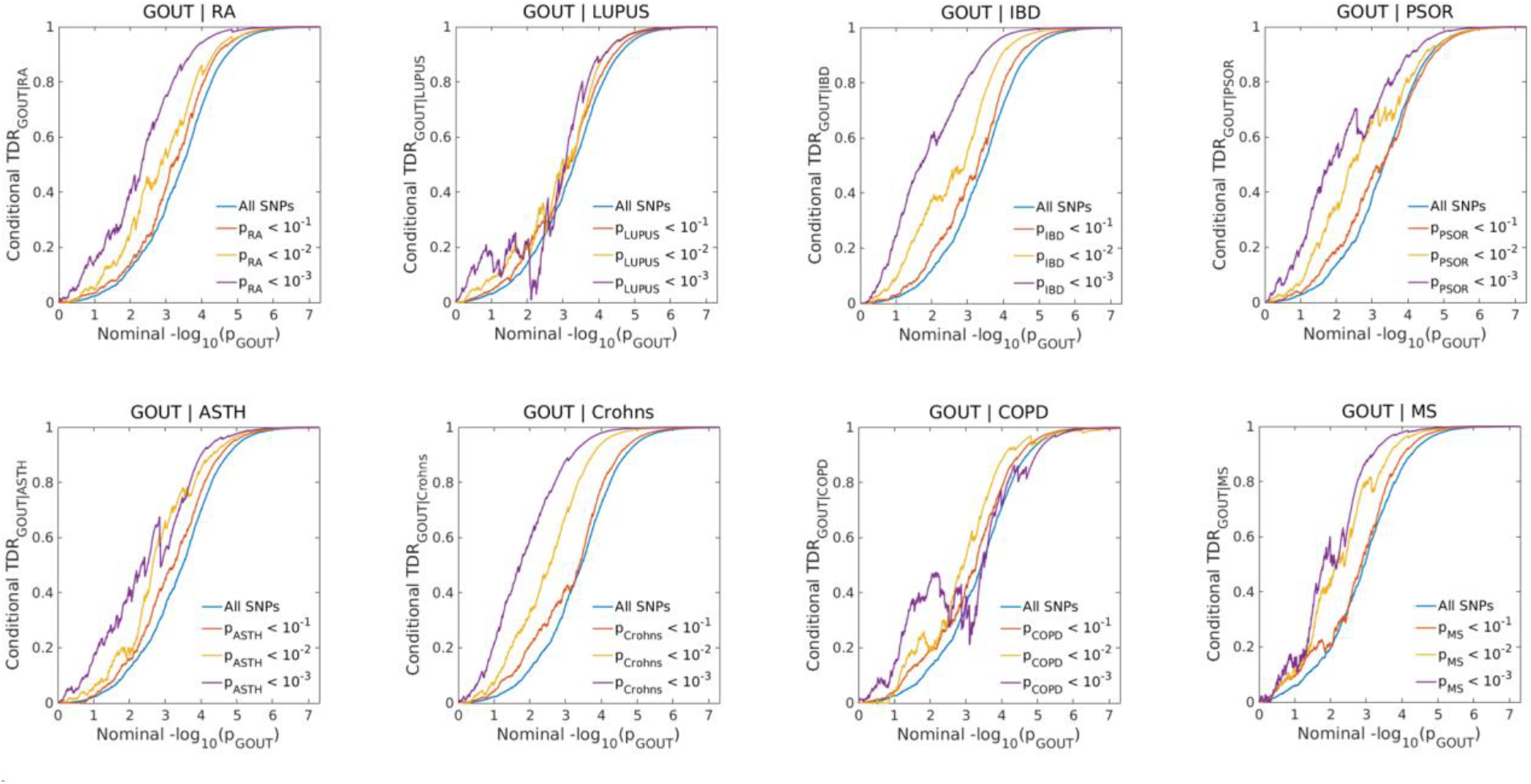
Conditional true discovery rate (TDR) plots showing the estimated proportion of true gout associations versus nominal –log_10_ P values, conditioned on different significance thresholds in eight immune-mediated disorders. Lines represent all SNPs (blue) and SNP subsets with increasing significance in the conditioning trait (orange, yellow, purple).

The conjFDR heatmaps (*Figure 3*) summarizes the joint significance landscape for each gout–immune disorder pair. In all comparisons, bright yellow regions in the lower right quadrant correspond to SNPs with strong associations in both traits, with the size and intensity of these regions reflecting the degree of pleiotropic sharing. Large, well-defined high-confidence regions were seen for Crohn’s disease, RA, IBD, psoriasis, and MS, whereas asthma, COPD, and SLE presented smaller but distinct overlap zones. The heatmap patterns align with both the TDR trajectories and the QQ plot deviations, collectively demonstrating that while the extent of genetic sharing differs among the eight immune-mediated disorders, pleiotropic enrichment is evident across all trait pairs examined. Taken together, these results demonstrate that gout shares a measurable and biologically meaningful proportion of its genetic architecture with a diverse range of immune-mediated diseases. The consistent patterns across all analyses underscore the robustness of these signals and indicate that the overlap is not driven by statistical artefacts but reflects genuine pleiotropic mechanisms. This broad distribution of loci across multiple chromosomes, coupled with trait-specific differences in enrichment magnitude, suggests the presence of both shared and disease-specific immune pathways that may contribute to gout pathogenesis.

**Figure 3.**
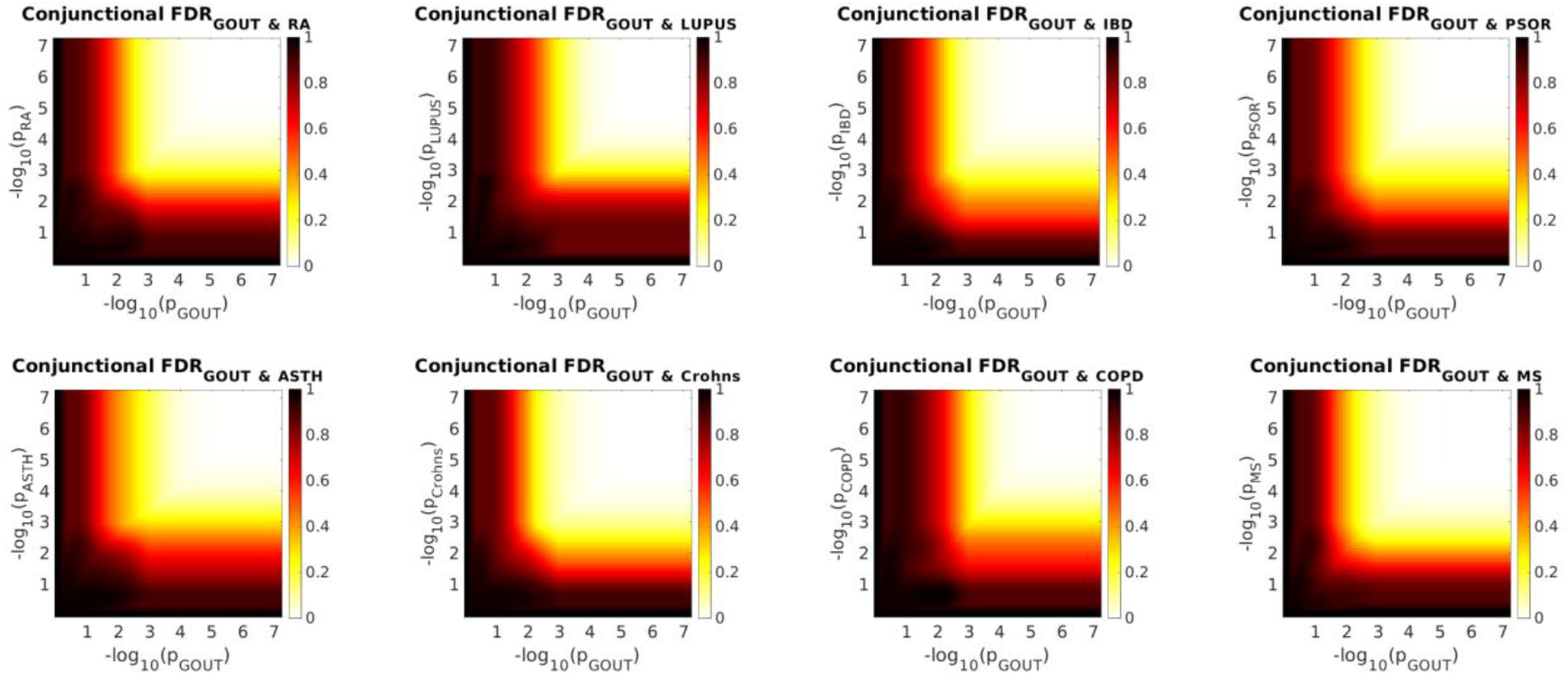
Conjunctional false discovery rate (conjFDR) heatmaps for gout and eight immune-mediated disorders. The x- and y-axes show –log_10_ P values for gout and the conditioning trait, respectively. Color scale indicates conjFDR values, with lighter colors representing stronger joint associations between the two traits.

**Figure 4.**
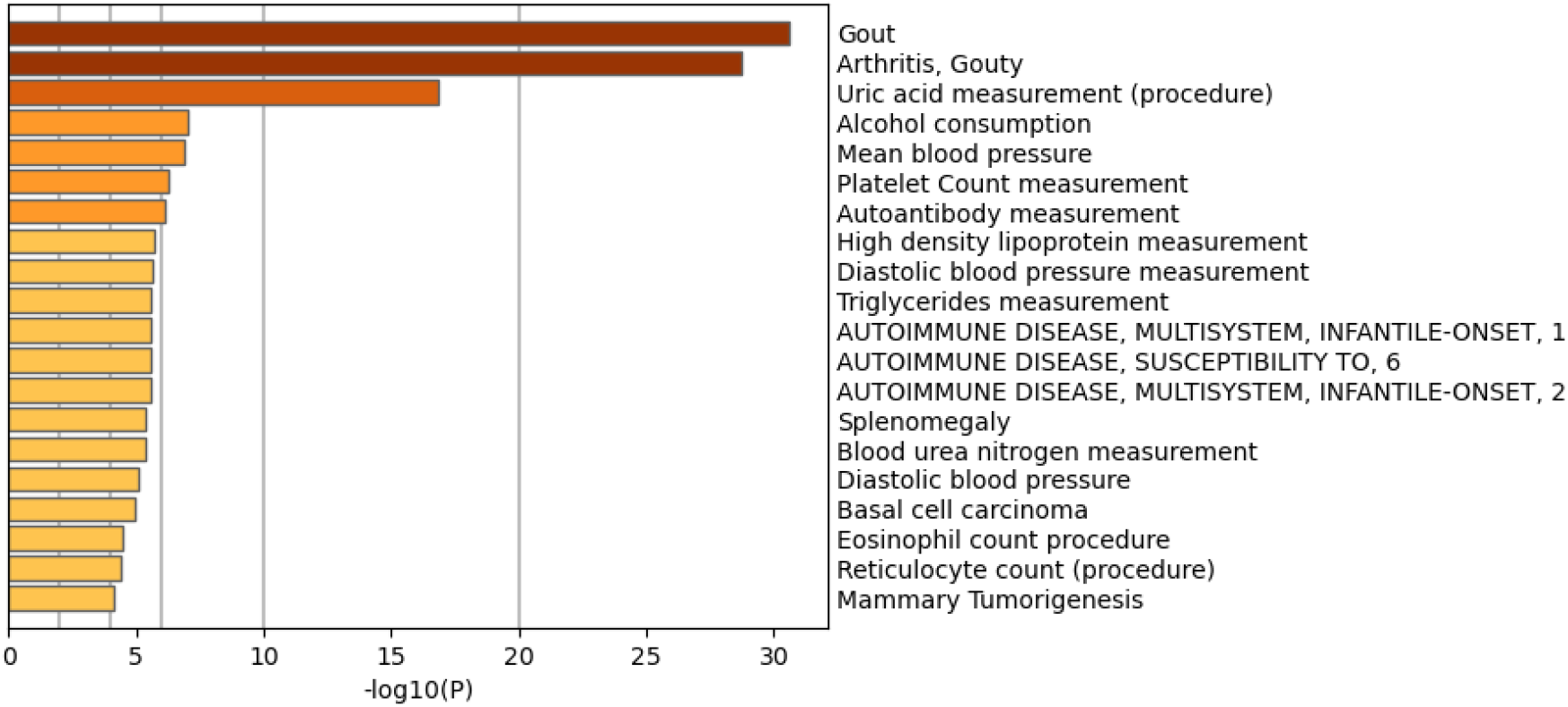
DisGeNET enrichment analysis of the 85 unique credible genes from the gout immune component. The x-axis shows –log_10_(p) values for associations between the gene set and curated disease or clinical measurement terms.

### Functional characterization of shared genetic loci between gout and immune-related traits

For each gout–immune disorder pair, the pleiotropic loci identified in the conjFDR analysis were individually processed through FUMA’s SNP2GENE pipeline, which applied positional overlap, eQTL colocalization, and promoter–capture Hi-C mapping strategies. Genes were retained as credible if they lay within 10 kb of a lead SNP and were supported by at least one functional criterion. This mapping was performed separately for each comparison, ensuring that per-disease gene lists were preserved to quantify gene-level sharing without losing resolution. Across all eight comparisons, this process yielded 152 credible gene assignments, with counts ranging from 1 in COPD and 5 in SLE to as many as 51 in Crohn’s disease and 42 in IBD, with intermediate numbers for RA (11), asthma (11), MS (17), and psoriasis (14). When these lists were combined and duplicates removed, the catalogue collapsed to 85 unique credible genes representing the entire gout immune component.

Despite the heterogeneous inputs, these 85 genes reduced to a compact pleiotropic core. Five genes— *ALDH2, HECTD4, MAPKAPK5, RP11-162P23*.*2*, and *SH2B3*—appeared in five of the eight disease comparisons, marking them as central hubs within the shared architecture (Table 1). A smaller group of other genes recurred across multiple traits, while the majority were observed in only one or two comparisons. These hub genes were located on several chromosomes, including chromosomes 2, 4, 6, 11, and 12, and importantly, no classical urate transporter genes (such as *SLC2A9* or *ABCG2*) were identified. This shows that the immune component identified is related to gout and independent of hyperuricemia. Additionally we identified 16 genes novel genes associated with gout shown in table 2.

**Table 1:** Top genes linked to ≥3 immune traits.

| Gene Symbol | Hits | Conditions |
| --- | --- | --- |
| *ALDH2* | 5 | Crohn's, IBD, LUPUS, PSOR, RA |
| *HECTD4* | 5 | Crohn's, IBD, LUPUS, PSOR, RA |
| *MAPKAPK5* | 5 | Crohn's, IBD, LUPUS, PSOR, RA |
| *RP11-162P23.2* | 5 | Crohn's, IBD, LUPUS, PSOR, RA |
| *SH2B3* | 5 | Crohn's, IBD, LUPUS, PSOR, RA |
| *ERBB3* | 3 | ASTH, COPD, PSOR |
| *KIAA1841* | 3 | Crohn's, IBD, MS |
| *OVOL1* | 3 | ASTH, MS, PSOR |
| *PUS10* | 3 | Crohn's, IBD, MS |
| *RNASEH2C* | 3 | ASTH, MS, PSOR |
| *USP34* | 3 | Crohn's, IBD, PSOR |

**Table 2:** Novel Genes associated with gout by leveraging shared genetic heritability with other immune disorders.

| **Gene Symbol** | **Hits** | **Conditions** |
| --- | --- | --- |
| *AHSA2* | 2 | Crohn's, IBD |
| *AMIGO3* | 2 | Crohn's, IBD |
| *BACH2* | 2 | Crohn's, IBD |
| *FAM212A* | 2 | Crohn's, IBD |
| *GBA* | 2 | Crohn's, IBD |
| *GMPPB* | 2 | Crohn's, IBD |
| *METTL10* | 2 | Crohn's, IBD |
| *MON1A* | 2 | Crohn's, IBD |
| *MST1R* | 2 | Crohn's, IBD |
| *RNF123* | 2 | Crohn's, IBD |
| *UBA7* | 2 | Crohn's, IBD |
| *KHK* | 1 | Crohn's |
| *PCNXL3* | 1 | MS |
| *PIP4K2C* | 1 | RA |
| *RP11-603J24.9* | 1 | ASTH |
| *TPH1* | 1 | Crohn's |

To investigate the functional context of the gout immune component, we used the set of 85 unique credible genes identified through conjFDR and FUMA mapping as input for gene set enrichment analysis. The immune component was assessed for enrichment using the GO and KEGG pathway databases to identify overrepresented biological processes, molecular functions, and signaling pathways. Using a stringent significance threshold of q-value (FDR-adjusted p-value) < 0.05, KEGG pathway analysis identified a focused set of immune-related processes, including T cell receptor signaling, B cell receptor signaling, antigen processing and presentation, NF-κB signaling, and Jak–STAT signaling, alongside multiple cytokine–cytokine receptor interaction terms. GO biological process enrichment converged on categories such as lymphocyte activation, leukocyte cell–cell adhesion, regulation of immune effector processes, and positive regulation of cytokine production. Together, these high-confidence pathways point to core adaptive and innate immune activation mechanisms, supporting that the gene set is indeed representative of the immune component of gout. The complete list of enriched pathways at this threshold is provided in Supplementary Table 1.

When applying a nominal p-value < 0.05 without multiple-testing correction — appropriate here given the exploratory aim and the prior enrichment of the gene set — a broader spectrum of potentially relevant pathways emerged. In KEGG, these included IL-6 signaling, IL-7 signaling, T helper cell differentiation, chemokine signaling, and Toll-like receptor signaling. GO biological processes expanded to cover mononuclear cell migration, interleukin-6 production, interleukin-7-mediated signaling, and positive regulation of T cell proliferation. While some of these nominal associations may represent false positives, many — notably IL-6 and IL-7 signaling — align closely with established models of gout inflammation, suggesting additional cytokine-driven mechanisms that could contribute to the shared genetic architecture between gout and other immune-mediated disorders. The complete list of nominally enriched pathways is provided in Supplementary Table 2.

Cross-referencing the 85 unique credible genes from the gout immune component against the DisGeNET database (Figure 5) identified significant associations with gout and gouty arthritis as the top-ranked traits, followed by clinical measurements such as uric acid levels, mean and diastolic blood pressure, platelet count, triglycerides, and high-density lipoprotein levels. Additional links were observed with alcohol consumption, autoantibody measurement, and several immune-mediated and multisystem disorders, as well as hematological traits including eosinophil and reticulocyte counts. The predominance of gout, arthritis, and immune-related conditions in the enrichment profile supports the interpretation that this gene set represents the immune component of gout rather than urate-handling pathways.

### Mendelian Randomization

To evaluate whether the genes identified through conjFDR and FUMA analyses play a causal role in the development of gout, we conducted a two-sample Mendelian Randomization (MR) analysis using blood-derived gene expression (eQTL) data. Among the 85 genes that constituted the immune component of gout, only 14 were found to exhibit significant causal relationships with gout in the MR framework. These genes represent a refined subset with functional relevance to disease etiology beyond genetic association alone (Table 3).

**Table 3:**
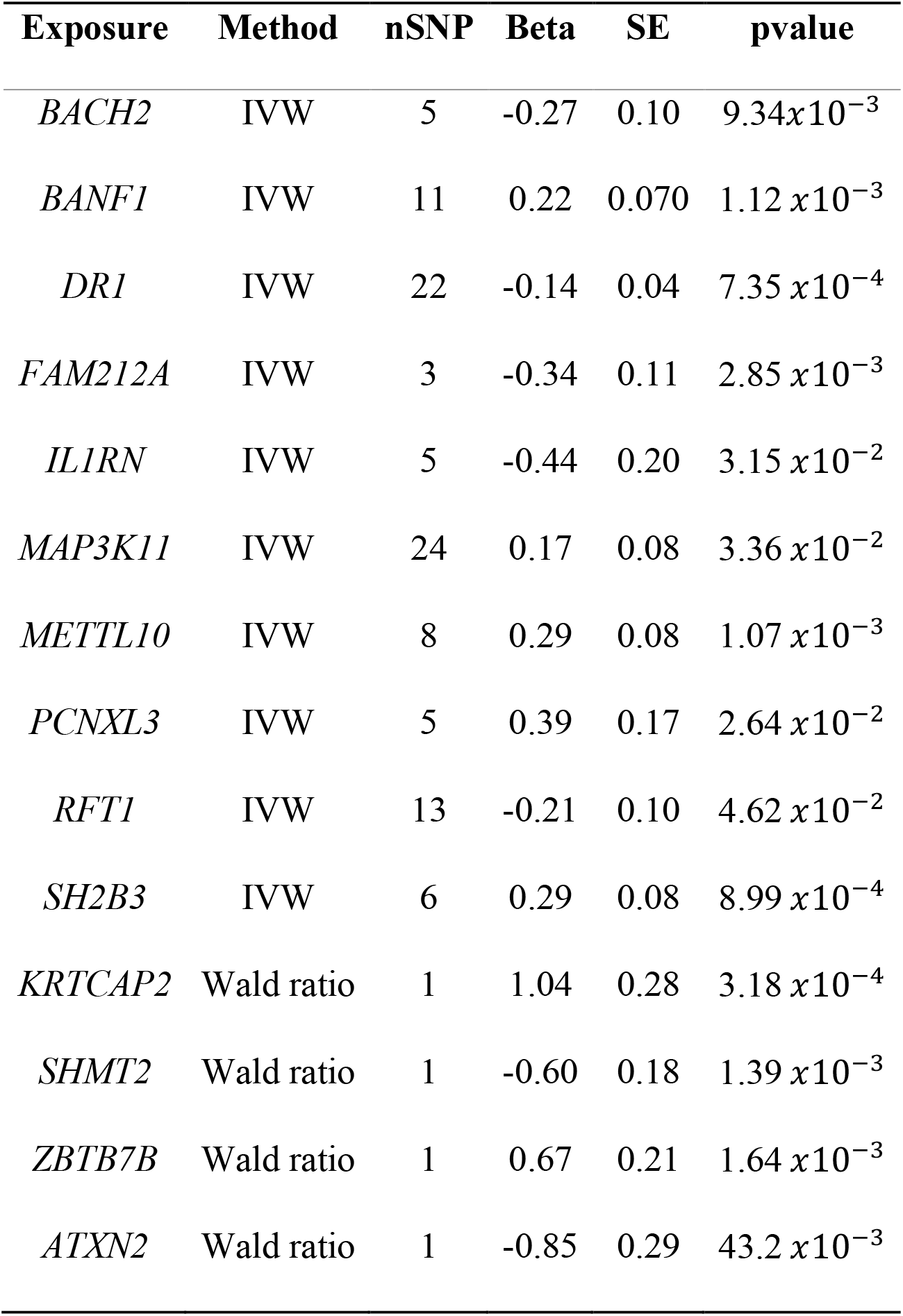
Mendelian-randomization summary: genetically predicted expression of the 85-gene immune component elevates gout risk.

**Table 4:** Therapeutic agents for gout based on the target genes.

| Drug | Target | Original Purpose | MoA | Source | Status |
| --- | --- | --- | --- | --- | --- |
| Pazopanib | *SH2B3* | Renal Cell Carcinoma | Inhibitor | CMap | Launched |
| CEP-1347 | *MAP3K11* | MLK Inhibitor | Inhibitor | DGIdb | Unknown |
| Candesartan | *SH2B3* | Hypertension | Unknown | DGIdb | Launched |
| URMC-099 | *MAP3K11* | HIV | Inhibitor | CMap | Preclinical |

Of the 14 genes with significant results, 10 demonstrated a positive association between increased gene expression and higher risk of gout, highlighting their potential as therapeutic targets for gene expression suppression or inhibition. Conversely, four genes displayed protective effects, where increased expression was associated with a decreased risk of gout. Notably, several genes with known immune regulatory roles, such as *SH2B3, IL1RN, DR1*, and *BACH2*, emerged with significant and biologically plausible MR estimates.

Five transcripts showed positive causal effects of higher expression on gout risk by IVW. The largest effect was for *PCNXL3* (β=0.39, SE=0.17, p=2.64 *x*10^−2^). Two signals had similar effect sizes but stronger significance: SH2B3 (β=0.29, SE=0.08, p=8.99 *x*10^−4^) and *METTL10* (β=0.29, SE=0.08, p=1.07 *x*10^−3^). *BANF1* also associated with increased risk (β=0.22, SE=0.07, p=1.12 *x*10^−3^), and *MAP3K11* showed a smaller but significant positive effect (β=0.17, SE=0.08, p=3.36 *x*10^−2^). Weighted-median (WM) estimates corroborated IVW for *SH2B3, METTL10, MAP3K11*, and *PCNXL3* (all WM p<0.05), but not for *BANF1* (WM p≈0.28). MR-Egger intercepts for these multi-SNP instruments provided no evidence of directional pleiotropy (all intercept p>0.05).

Five transcripts exhibited negative effects, consistent with protection at higher expression. The largest magnitude was *IL1RN* (β=−0.44, SE=0.20, p=3.15 *x*10^−2^), followed by *FAM212A* (β=−0.34, SE=0.11, p=2.85 *x*10^−3^) and *BACH2* (β=−0.27, SE=0.10, p=9.34*x*10^−3^). *RFT1* showed a modest protective effect (β=−0.21, SE=0.10, p=4.62 *x*10^−2^). Notably, *DR1* had the smallest effect size but the strongest statistical support among negatives (β=−0.14, SE=0.04, p=7.35 *x*10^−4^). WM supported *BACH2, FAM212A*, and *DR1* (all WM p<0.05), was borderline for RFT1 (WM p≈0.067), and did not replicate *IL1RN* (WM p≈0.34). MR-Egger intercepts were again non-significant across these traits (all intercept p>0.05; range ≈0.08–0.72), arguing against directional pleiotropy.

Three exposures were instrumented by a single eQTL and analyzed with the Wald ratio. The largest effect was *KRTCAP2* (β=1.04, SE=0.28, p=3.18 *x*10^−4^), followed by *ZBTB7B* (β=0.67, SE=0.21, p=1.64 *x*10^−3^); both indicate higher expression increases risk. In contrast, *SHMT2* suggested protection (β=−0.60, SE=0.18, p=1.39 *x*10^−3^). For these one-SNP instruments, WM and MR-Egger are not applicable.

### Drug repurposing

From the 85 genes constituting the immune component of gout, Mendelian Randomization identified 14 with a causal role in disease risk. Among these, five genes—*BANF1, MAP3K11, PCNXL3, SH2B3*, and *METTL10*—showed upregulation and were prioritized as therapeutic targets. These were input into DGIdb and ConnectivityMap, revealing four existing compounds with potential relevance to gout therapy: pazopanib (*SH2B3* inhibitor), CEP-1347 and URMC-099 (*MAP3K11* inhibitors), and candesartan (targeting *SH2B3*, though with an unclear mechanism of action).

## DISCUSSION

The primary objective of this study was to isolate and characterize the immune-specific component of gout by leveraging data from eight well-established immune-mediated disorders in a conjFDR framework. We identified 85 unique credible genes representing the shared immune architecture between gout and autoimmune or inflammatory conditions. Several hub genes, including *ALDH2, SH2B3, MAPKAPK5, KIAA1841*, and *OVOL1*, were shared between gout and five immune disorders, suggesting conserved effector modules contributing to common inflammatory pathways. Gene Ontology enrichment revealed significant overrepresentation of immune-related processes such as leukocyte activation, neutrophil degranulation, cytokine signaling, and NF-κB regulation—consistent with an inflammatory cascade triggered by monosodium urate crystals rather than directly modulating the NLRP3 inflammasome, a finding that is supported by the most recent GWAS of gout by Major et al have identified loci that are involved in cytokines and regulation of the NLRP3 inflammasome response in gout (Major et al., 2024). DisGeNET enrichment analysis of the immune component supports our conclusion that the gene set proposed can be termed the immune component of gout, as the top two terms—Gout and Arthritis, gouty—confirm that the signal concentrates on the disease itself. Furthermore, almost all gout patients undergo uric acid measurement procedures, so the term appearing third on the list of conditions is understandable. Alcohol consumption appearing fourth is not surprising as the relationship between alcohol consumption and gout has a long history, through most of which alcohol consumption has been seen as a cause of developing gout, but more recently Mendelian randomization has shown that the relationship is actually in the opposite direction, meaning individuals with a genetic predisposition towards gout tend to have a genetic predisposition towards drinking more. Importantly, the remainder of the profile is enriched for immune and hematologic readouts—autoantibody measurement, eosinophil and reticulocyte counts, splenomegaly—and formal autoimmune disease terms, collectively reinforcing that this gene set captures an immunogenetic component of gout rather than general urate-handling biology. Finally, the absence of the immune diseases used to characterize the immune component of gout from among the leading terms indicates that enrichment is specific to gout and not driven by those other diseases.

Cross-referencing this 85-gene panel against all gout GWAS published through December 2024 revealed three tiers of evidence supporting its validity. Approximately 78% (66 genes) reside within loci previously associated with gout, while three genes (*ERBB3, MERTK, TMEM87B*; ∼4%) were identified in the 2024 trans-ancestry GWAS by major et al for the first time, validating our approach (Major et al., 2024). Notably, 19% (16 genes) represent novel candidates not previously linked to gout. Among these, five genes—*BACH2, MST1R, RNF123, UBA7*, and *PIP4K2C*—exhibit well-established immune functions based on prior functional genomics and immunology studies. *BACH2* encodes a transcription factor that maintains T cells in a naïve state, regulates Th2 differentiation, and supports T-regulatory cell function, thereby ensuring adaptive immune balance (Liu & Liu, 2022; Tsukumo et al., 2013; Zwick, Vo, Shim, Reijonen, & Do, 2024), additionally results of our MR analysis show that *BACH2* causally affects the risk of developing gout. *MST1R* (RON receptor tyrosine kinase) is expressed on tissue-resident macrophages and epithelial cells, where it inhibits classical macrophage activation, promotes alternative activation linked to wound healing, and regulates cytokine responses such as IFN-γ production (Wilson et al., 2008). *RNF123* mediates ubiquitination and degradation of *SOCS1*, thereby regulating type I interferon production and antiviral immunity (Huang et al., 2023). *UBA7* encodes the E1 activating enzyme for ISG15, with structural studies confirming its role in interferon-induced ISG15 activation, a critical antiviral defense mechanism (Afsar et al., 2023). *PIP4K2C* deletion in murine models results in hyperactivation of the immune system, underscoring its role in restraining T-cell activation and cytokine responses (Shim et al., 2016). The presence of these genes within the novel component of our gout immune signature reinforces the conclusion that our pleiotropy-guided approach has identified biologically plausible targets within immune regulatory networks, extending beyond the urate-handling genes typically recovered in gout GWAS.

From a drug discovery perspective, approximately one-third of our prioritized genes encode enzymes or receptors with tractable binding sites. Advances in structure-guided ligand design, high-throughput phenotypic screening, and artificial intelligence-driven cheminformatics could accelerate the development of first-in-class therapeutics targeting gout’s immune component. To prioritize druggable targets, we performed two-sample Mendelian randomization using blood eQTL data as exposure identifying 14 genes whose expression in blood significantly influenced gout risk. Among these, *IL1RN, MAP3K11*, and *SH2B3* stand out due to existing pharmacological modulators. Mechanistically, *MAP3K11* drives inflammatory amplification via the *JNK* and p38 *MAPK* pathways; its inhibition could blunt both acute and chronic inflammation (Bachstetter & Van Eldik, 2010; Kumar et al., 2020). *SH2B3* regulates JAK/STAT signaling, and although its inhibition may dampen cytokine production, potential risks include broader immune suppression (Morris, Butler, Perkins, Kershaw, & Babon, 2021). Reduced *IL1RN* expression was causally linked to gout risk, supporting *IL-1* blockade as a therapeutic strategy—a concept validated by prior trials and reinforced by recent Phase 3 data for Firsekibart in China (Xue et al., 2025). *IL-1* inhibition serves as a proof-of-concept (Janssen et al., 2019), but additional strategies such as *MAP3K11* blockade, *MST1R* or *PIP4K2C* inhibition etc., could provide complementary benefits while minimizing infection risks. This study strengthens the evidence to support immune-mediated mechanisms in gout, identifying *IL1RN, MAP3K11*, and *SH2B3* as causal and druggable targets. These findings, corroborated by recent GWAS and multi-omics studies, highlight the potential of immune-targeted therapies to complement urate-lowering strategies.

This is the first study of its kind to attempt to characterize the genetic immune component of gout and broadened our understanding of gout as an immune disorder by both proposing novel genes and possible targets within immune component. However, the greatest limitation of using the conjFDR method to determine the immune component of gout is that it only identifies shared inheritance and may inadvertently identify genes that are shared but not due to shared immune components. Although the use of immunotherapy in the treatment of gout is slowly gathering pace and being prescribed more often by clinicians, due to the complex nature of the immune regulation further research into the immune nature of gout is required to understand how to best integrate it into clinical practice.

